# Based on the identification of tumor antigens and immune subtypes in renal clear cell carcinoma for mRNA vaccine development

**DOI:** 10.1101/2021.07.04.451033

**Authors:** Shichao Zhang, Yu Xiong, Shijing Kang, Chengju Mao, Yue Wang, Zhu Zeng, Jian Peng, Yan Ouyang

**Author notes:** Corresponding authors: A/Prof. Jian Peng, Key Laboratory of Infection Immunity and Antibody Engineering of Guizhou Province, School of Biology and Engineering & School of Basic Medical Sciences, Guizhou Medical University, Guiyang 550025, China. A/Prof. Yan Ouyang, Key Laboratory of Infection Immunity and Antibody Engineering of Guizhou Province, School of Biology and Engineering & School of Basic Medical Sciences, Guizhou Medical University, Guiyang 550025, China. Prof. Zhu Zeng, Key Laboratory of Infection Immunity and Antibody Engineering of Guizhou Province, School of Biology and Engineering & School of Basic Medical Sciences, Guizhou Medical University, Guiyang 550025, China.

## Abstract

**Background:** Cancer vaccine based on mRNA is considered as a promising strategy and has become a new hot spot in cancer immunotherapy. However, its application to KIRC is not clear. A growing body of research has shown that immunotyping can reflect the comprehensive immune status and immune microenvironment of tumor, which is closely related to treatment response and vaccination potential. The aim of this study was to identify the potential antigens of KIRC for the development of anti-KIRC mRNA vaccines, and to further differentiate the immune subtypes of KIRC to construct an immune landscape for the selection of appropriate patients for vaccination.

**Methods:** Gene expression profiles and corresponding clinical information of 265 KIRC patients and RNA-seq data of 539 KIRC patients were retrieved from were collected from GEO and TCGA. cBioPortal was used to visualize and compare genetic alterations, while GEPIA2 was used to calculate the prognostic index of selected antigens. The relationship between the infiltration of antigen presenting cells and the expression of the identified antigen was visualized with TIMER, and consensus clustering analysis was used to determine the immune subtypes. Finally, the immune landscape of KIRC is visualized through the dimensionality reduction analysis based on graph learning.

**Results:** Two tumor antigens associated with prognostic and antigen-presenting infiltrating cells were identified in KIRC, including LRP2, and DOCK8. KIRC patients were classified into six immune subtypes based on different molecular, cellular, and clinical characteristics. Patients with IS5 and IS6 tumors had an immune “hot” and immunosuppressive phenotype, which was associated with better survival compared to other subtypes, whereas patients with IS1-4 tumors had an immune “cold” phenotype, which was associated with a higher tumor mutation burden. In addition, the expression of immune checkpoints and immunogenic cell death modulators differed significantly in different immunosubtypes of tumors. Finally, the immune landscape of KIRC shows a high degree of heterogeneity across patients.

**Conclusions:** LRP2 and FEM2 are potential KIRC antigens for mRNA vaccine development, and patients with immune subtypes IS1-4 are suitable for vaccination.

## Introduction

Renal cell carcinoma (RCC) affects over 400,000 individuals worldwide per year [1]. It predominantly affects individuals over 60 years of age, and two-thirds of patients are male [1]. It is anticipated that 73,750 people would be newly diagnosed with kidney cancer and 14,830 patients are likely to die because of the disease in the USA in 2020 [2]. Several subvariants of RCC exist, with approximately 70% of individuals being diagnosed with Kidney renal clear cell carcinoma (KIRC) [3]. Although KIRC can be successfully treated with surgical or ablative strategies if identified early, up to a third of patients will present with or develop metastases[4]. Metastatic KIRC is almost uniformly lethal and is biologically distinct from non-metastatic disease. Surgery excision remains the primary therapy for KIRC due to the growing resistance to radiotherapy and chemotherapy [5]. Much worse, the prognosis of KIRC patients tends to be poor, especially for patients in an advanced stage. The five-year overall survival rate of stage IV patients is less than 10% [5].

Previous studies have revealed that the presence of classable diversity in the immunological signature of tumor tissue-infiltrating lymphocytes is significantly linked to the survival of KIRC patients [6, 7], and immunotherapy has been suggested as the treatment for metastatic KIRC [8, 9]. To date, cancer immunotherapy has achieved considerable success in combatting several malignancies [10]. Following immune checkpoint inhibitors targeting programmed cell death protein 1 and its ligand 1, the mRNA-cancer vaccine has become increasingly attractive to scientists and oncologists and could be a hotspot in cancer immunotherapy [11, 12]. In fact, attempts at mRNA-based therapies were not common until the 2000s due to the instability of mRNA and the associated excessive inflammatory response. However, technological breakthroughs in recent years, including the addition of modified nucleosides, purification of engineered in vitro transcription (IVT) mRNAs, optimization of coding sequences, and development of effective delivery materials that enable mRNAs to carry tumor antigens in an optimal form, have changed this situation [12, 13]. For instance, mRNA sequences can be easily modified to produce any protein as needed, while traditional peptide vaccines require complex genetic analysis of cancer. This greatly improves the production efficiency and cost of the vaccine, and shortens the empty window of treatment for cancer patients. Importantly, the safety of mRNA can be improved by regulating the half-life of mRNA through delivery system or RNA sequence modification. Moreover, compared with DNA vaccines, mRNA vaccines do not have the risk of gene integration, insertion mutations or gene deletions caused by DNA-type vaccines. It has been found that the self-adjuvant properties of mRNA can increase its immunogenicity in vivo and induce a strong and sustained immune response [10, 12, 14]. Preclinical and clinical models have demonstrated that vaccines encoding tumor-specific antigens promote anti-tumor immunity and prevent a wide range of tumors, including melanoma, hepatocellular carcinoma, colorectal cancer, gastrointestinal cancer, and pancreatic adenocarcinoma [15–17].

However, KIRC still lacks an effective mRNA vaccine because it remains challenging to isolate effective antigens for use against KIRC from the thousands of mutant candidate genes in tumor cells [18]. Moreover, due to the high heterogeneity of tumor cells and their complex tumor immune microenvironment (TIME), only a small percentage of KIRC patients are likely to benefit from mRNA vaccines [19–22]. Therefore, patient stratification based on tumor biological subtypes can be used to identify patients suitable for vaccination, thereby improving the accuracy of diagnosis and treatment. Previous KIRC classifications, which are based on specific molecular patterns, have focused on the intrinsic molecular profile of tumor cells, including gene amplification, copy number alterations, and signaling pathway dysregulation, with a few focusing on tumor-infiltrating immune cells.[4, 23, 24]. However, from the point of view of the interaction between tumor cells and the body’s immune system, these traditional methods are not sufficient to screen candidate genes for mRNA vaccines. In contrast, stratification of immune gene expression profiles in patients allows identification of patients suitable for mRNA vaccination from immunologically heterogeneous populations[25, 26].

This study aimed to identify novel KIRC antigens for developing mRNA vaccines, and map the immune landscape of KIRC to select suitable patients for vaccination.

Two candidate genes associated with survival and antigen-presenting cell (APC) infiltration were identified from the pool of overexpressed and mutated genes in KIRC. 6 robust immune subtypes and 8 functional modules of KIRC were identified by immune-associated gene clustering, each corresponding to different clinical, molecular, and cellular characteristics, and these subtypes were validated in an independent cohort. Finally, the immune profile of KIRC is defined by analyzing the distribution of relevant genetic signals in each patient. Our findings define the complex TIME of each KIRC patient and provide a theoretical basis for the selection of mRNA vaccine targets and the screening of suitable patients for vaccination.

## Methods and materials

### Data extraction

The normalized gene expression and clinical follow-up data of 265 KIRC patients, and RNA-seq data of 539 KIRC patients were retrieved from Gene Expression Omnibus (GEO, https://www.ncbi.nlm.nih.gov/geo) and The Cancer Genome Atlas (TCGA, https://www.cancer.gov/tcga) respectively through GDC API. A total of 2598 immune-related genes were extracted from the discovery and validation cohort, including single immune cell-specific, co-stimulatory and co-inhibitory molecules, cytokines and cytokine receptors, antigen processing and presentation, and other immune-related genes.

### Data preprocessing

To process the GEO data, the microarray probes with null gene test value were first removed and the remaining probes were mapped to human genes. Among the 2108 expressed genes, 782 immune cell-related genes were retained. In TCGA tumor samples, those lacking clinical information and normal tissue data were first excluded, and then 50% of the sample genes with 0 FPKM in > 50% were excluded. Finally, 733 immune cell-related genes with log2 (FPKM +1) fold-change FPKM were included for the subsequent analysis.

### GEPIA analysis

Differential gene expression and patient survival data were integrated using Gene Expression Profiling Interactive Analysis [27] (GEPIA, http://gepia2.cancer-pku.cn, version 2). ANOVA was used to identify the differentially expressed genes with |log2FC| values > 1 and q values < 0.01. Kaplan-Meier method with a 50% (Median) cutoff was used to evaluate Overall Survival (OS) and relapse-free Survival (RFS), and the log rank test was applied to compare them. In addition, the Cox proportional risk regression model was used to calculate the hazard ratio. Parameter Settings were consistent in each analysis, and no adjustments were made to any p values. *P* < 0.05 were considered statistically significant.

### cBioPortal analysis

The raw RNA-Seq data from TCGA, ICGC and other databases were integrated with the cBio Cancer Genomics Portal [28] (cBioPortal, http://www.cbioportal.org) to compare the genetic changes of KIRC. *P* < 0.05 were considered statistically significant.

### TIMER analysis

The relationship between the abundance of tumor immune infiltrating cells (TIICs) and KIRC-related genes was analyzed and visualized using Tumor Immune Estimation Resource [29] (TIMER, https://cistrome.shinyapps.io/timer/) through analysis modules including gene expression, somatic mutations, clinical outcomes, and somatic copy number alteration. Purity Adjustment was selected using Spearman’s correlation analysis. *P* < 0.05 were considered statistically significant.

### Discovery and validation of the immune subtypes

Cluster analysis was performed on the expression profiles of 2598 immune-related genes, and a consistency matrix was constructed to identify the corresponding immune subtypes and gene modules. The partition around medoids algorithm with the “1-Pearson correlation” distance metric was applied, and 500 bootstraps were performed each involving 80% patients in the discovery TCGA cohort. The range of cluster sets ranged from 2 to 10, and the optimal partition was obtained by evaluating the consensus matrix and consensus cumulative distribution function. The obtained immune subtypes were then validated in an independent GEO cohort using the same Settings. The consistency of immune subtypes between the discovery cohort and the validation cohort was quantified by calculating the intra-group proportion and Pearson correlation in the centroids of gene module scores.

### Immune-related molecular and cellular features

The “clusterProfiler” package was used for enrichment analysis of biological processes to explore signaling pathways associated with immune-related molecular and cellular characteristics [30]. The relationship between 56 immune-related molecular and cellular characteristics or immune subtypes was evaluated, and the composition of immune cells in tumor tissues deduced by the CIBERSORT algorithm was analyzed [31].

### Gene co-expression network

The R software package WGCNA was used to identify the immunorelated gene co-expression modules. Single-sample GSEA (SSGSEA) was used to calculate the immunoenrichment score for each sample to measure the coordinated up-regulation or down-regulation of genes within a sample.

### Construction of immune landscape

The reduce Dimension function of Monocle package with a Gaussian distribution was used to perform dimensionality reduction analysis based on graph learning to visualize the distribution of immune subtypes among individual patients. The maximum number of components was set to 2, and the discriminative dimensionality reduction with trees was used. Finally, the visualization of immune landscape was accomplished by using the functional plot cell trajectories of the color-coded immune subtypes.

## Results

### Identification of KIRC potential antigens

To identify potential antigens for KIRC, 1633 overexpressed genes that may encode tumor-associated antigens were detected by screening for aberrantly expressed genes (Figure 1A). After analyzing the altered genome fraction and the number of mutations in a single sample, 9843 mutated genes that may encode tumor-specific antigens were screened (Figure 1B, C). Mutational analysis showed that von Hippel-Lindau (VHL), and BRG1-Associated Factor 180 (PBRM1) were the most frequently mutated genes in terms of both altered genome fraction and mutation counts (Figure 1D, E). Most notably, in addition to VHL, PBRM1 in top 10 candidates with altered genome fractions, titin (TTN), collagen type XXIV alpha 1 chain (COL24A1), synergin gamma (SYNRG), alpha-2-macroglobulin like 1 (A2ML1), ADNP homeobox 2(ADNP2), adenosylhomocysteinase like 1 (AHCYL1), ALG10 alpha-1,2-glucosyltransferase B (ALG10B), as well as angiopoietin 1 (ANGPT1), all have the same mutation frequency, indicating the potential genomic interaction among them (Figure 1D). High mutation counts were observed in mucin 16 (MUC16), ArfGAP with RhoGAP domain, ankyrin repeat and PH domain 3 (ARAP3), teashirt zinc finger homeobox 3 (TSHZ3), dynein axonemal heavy chain 3(DNAH3), fibrillin 2 (FBN2), heme binding protein 1(HEBP1), SSX family member 3 (SSX3), and ATP binding cassette subfamily A member 6 (ABCA6) (Figure 1E). Overall, 684 overexpressed and frequently mutated tumor specific genes were identified.

**Figure 1.**
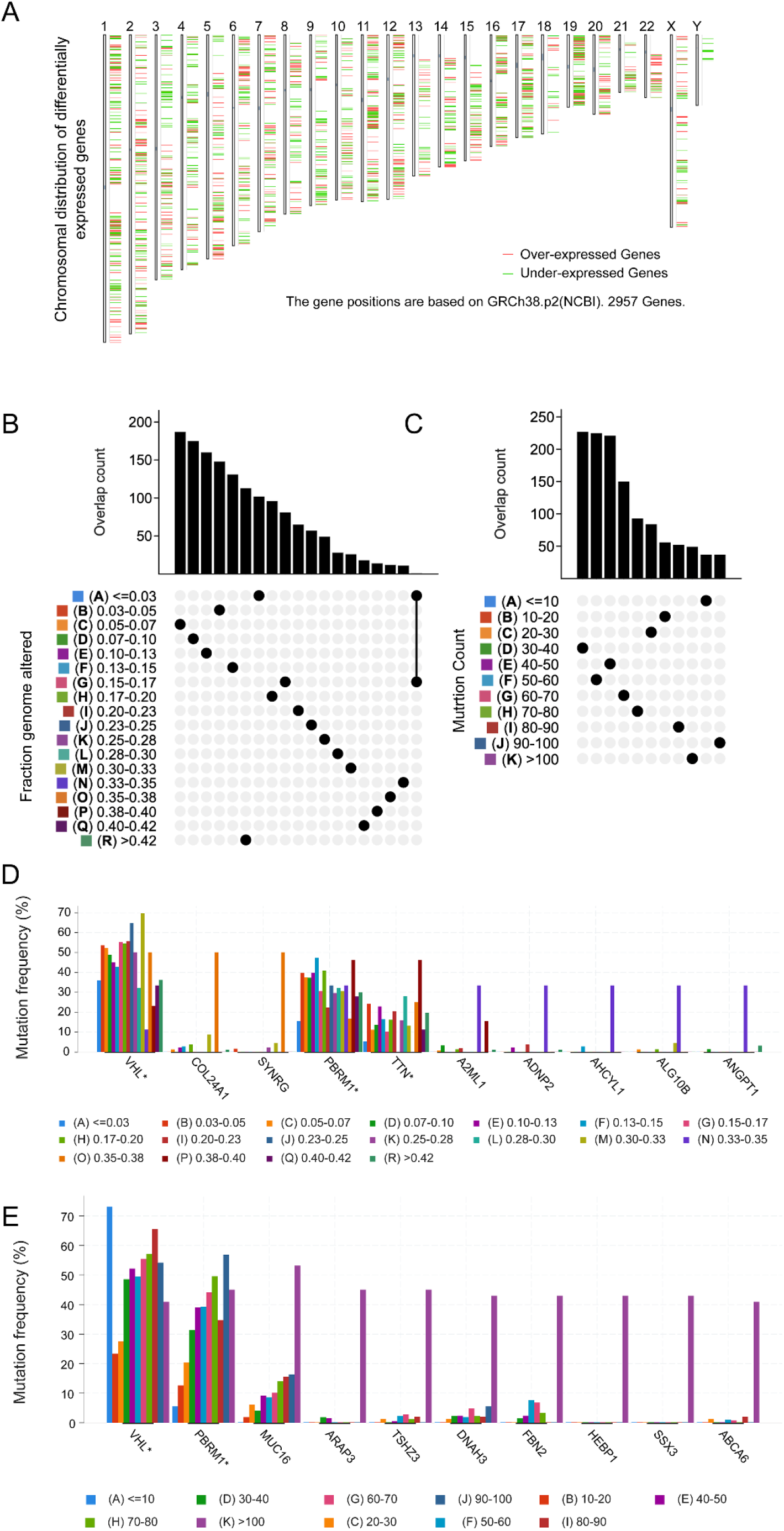
Identification of potential tumor antigens of KIRC. (A) Identification of potential tumor-associated antigens of KIRC. Chromosomal distribution of up- and down-regulated genes in KIRC as indicated. (B-E) Identification of potential tumor-specific antigens of KIRC. Samples overlapping in altered genome fraction (B) and mutation count groups (C), genes with highest frequency in altered genome fraction (D) and mutation count groups (E).

### Identification of tumor antigens associated with KIRC prognosis and antigen presenting cells

From the above genes, tumor genes related to patient prognosis were next screened as potential candidate antigens for developing mRNA vaccines. Eight genes were found to be closely related to OS of KIRC patients, and 2 of them were significantly related to RFS (Figure 2A). As shown in the Figure 2B-C, patients overexpressing LDL receptor related protein 2 (LRP2) in the tumor tissues had significantly longer survival compared to the LRP2^low^ group. Likewise, high expression levels of dedicator of cytokinesis 8 (DOCK8) was also associated with better prognosis (Figure 2D, E). In summary, 2 candidate genes were identified as having fundamental roles in the occurrence and progression of KIRC. In addition, high expression levels of LRP2 and DOCK8 were significantly associated with increased tumor infiltration of macrophages, DCs and B cells (Figure 3A, B). These results suggest that the identified tumor antigens may be directly processed by APC and presented to T cells, which are recognized by B cells to trigger an immune response, and thus are promising candidates for the development of an mRNA vaccine against KIRC.

**Figure 2.**
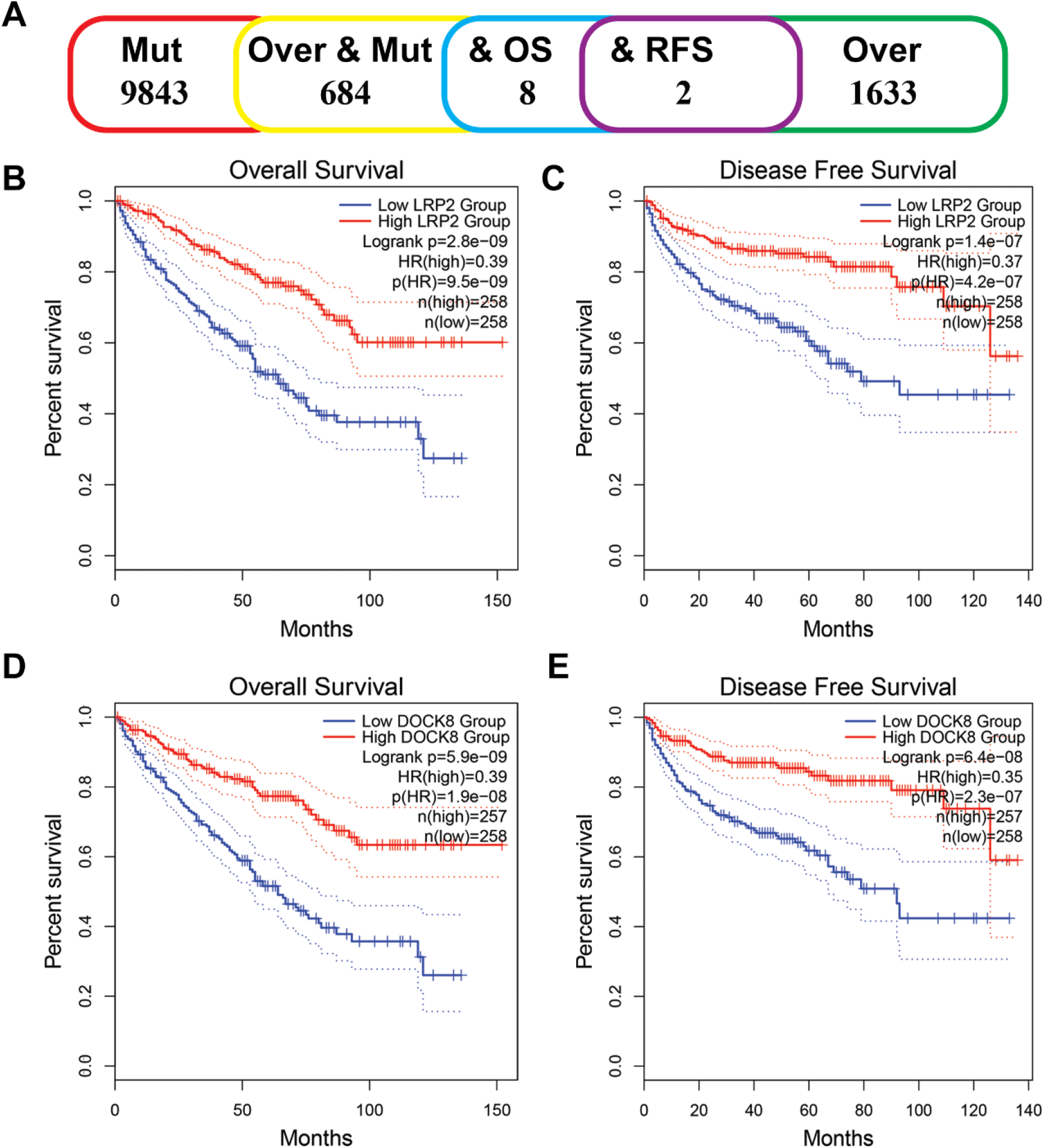
Identification of tumor antigens associated with KIRC prognosis. (A) Potential tumor antigens (total 139) with high expression and mutation in KIRC, and significant association with OS and RFS (total 2 candidates). Kaplan-Meier curves showing OS of KIRC patients stratified according to LRP2 (B), and DOCK8 (D) expression levels. Kaplan-Meier curves showing RFS of KIRC patients stratified according to LRP2 (C) and DOCK8 (e) expression levels.

**Figure 3.**
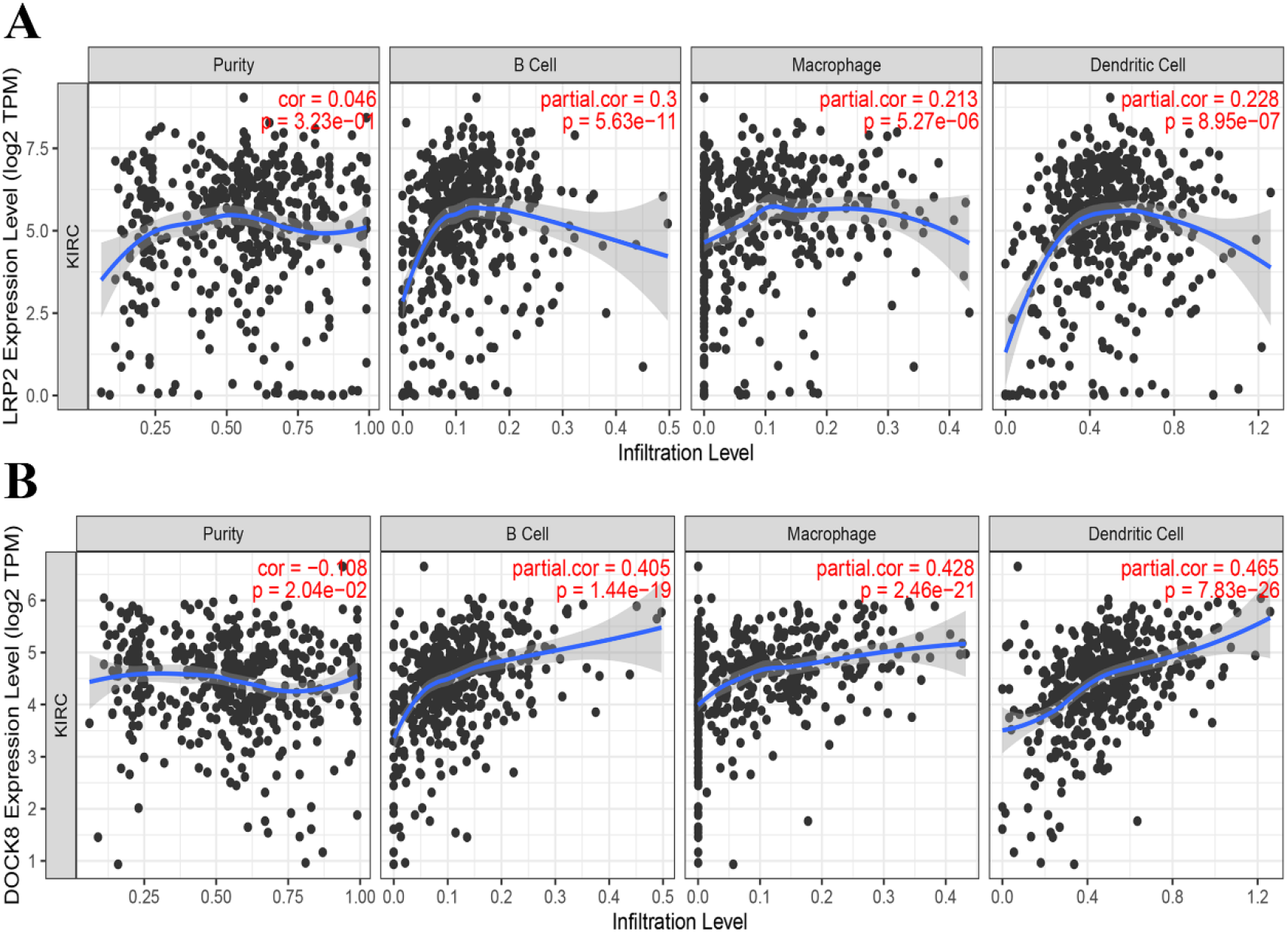
Identification of APCs-associated tumor antigens. Correlation between the expression levels of LRP2 (A), DOCK8 (B), and the infiltration of B cells, macrophages and dendritic cells in KIRC tumors

### Identification of potential immune subtypes of KIRC

Since the tumor immune response varies greatly among patients with different tumors, immunotyping can be used to reflect the immune status of the tumors and their microenvironment, thus helping to identify suitable patients for inoculation. Therefore, consensus clustering was constructed by analyzing the expression profiles of 2598 immune-related genes from 539 KIRC samples in the TCGA database. According to their cumulative distribution function and functional delta area, immunorelated genes showed stable cluster when k = 6 (Figure 4A and B), and 6 immune subtypes were obtained, respectively designated as IS1-IS6 (Figure 4C). IS5 and IS6 were associated with better prognosis, whereas IS4 had the poorest survival probability (Figure 4D). Subtype distribution across different tumor stages and grades indicated that IS5 and IS6 were significantly associated with lower tumor grade (Figure 4E), earlier clinical stage (Figure 4F) and T-stage (Figure 4G). In conclusion, immunotyping can be used to predict the prognosis of KIRC patients, and its predictive accuracy is superior to that of conventional grading and staging, which is consistent across cohorts.

**Figure 4.**
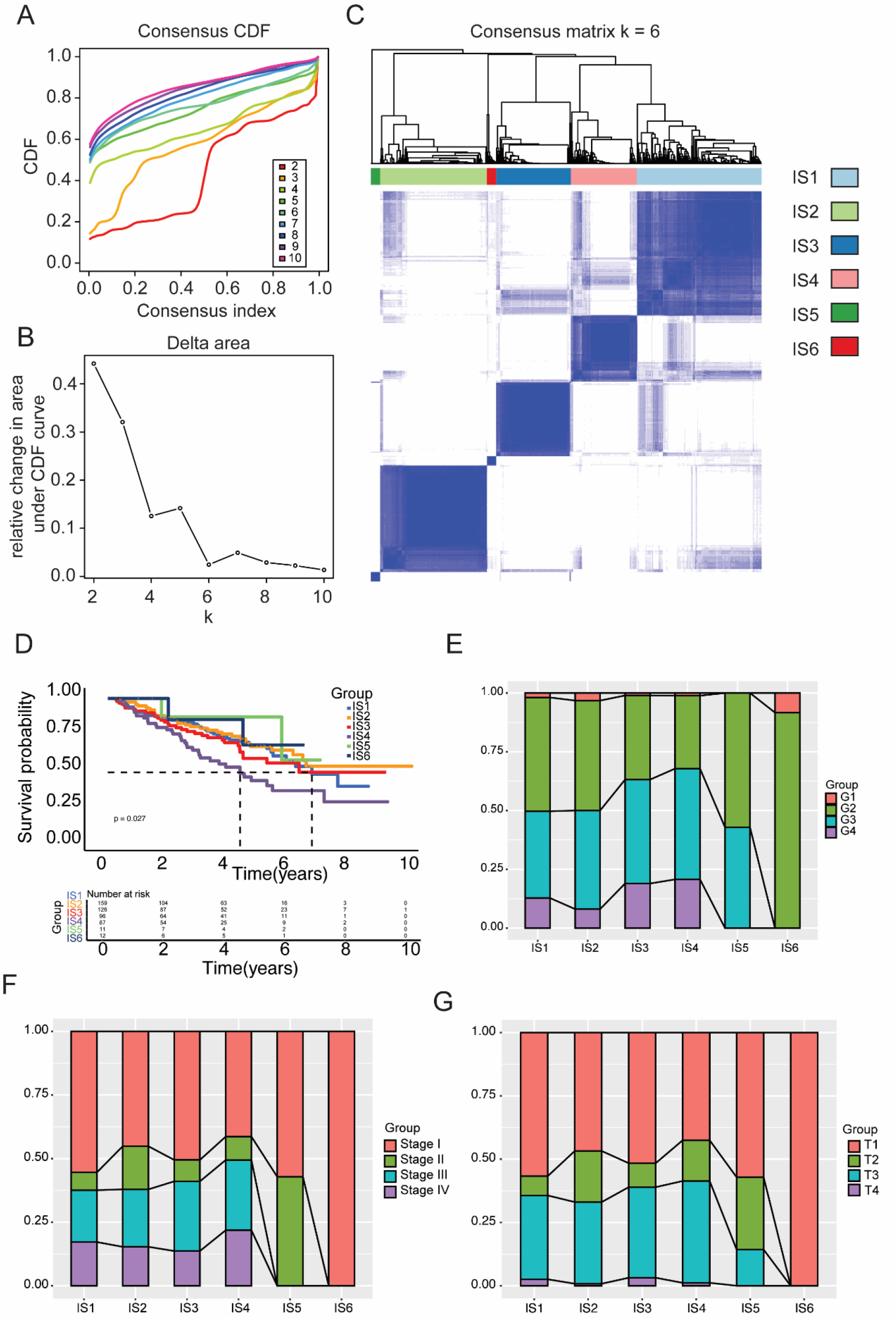
Identification of potential immune subtypes of KIRC. (A) Cumulative distribution function curve and delta area of immune-related genes in TCGA cohort (B). (C) Sample clustering heat map. D Kaplan-Meier curves showing OS of KIRC immune subtypes in TCGA cohort. (E, F, G) Distribution of IS1-IS6 across KIRC grades (E), stages (F), and Tumor stages of TNM (G) in TCGA cohort.

### The correlation of immune subtypes with mutational status

Previous Study has shown that higher tumor mutation burden (TMB) and somatic mutation rates in cancer patients are associated with stronger anti-cancer immunity [32]. Therefore, TMB and mutations were calculated for each patient with the mutect2-processed mutation dataset of TCGA, and the same analysis was performed for all immune subtypes. As shown in Figure 5A, IS5 and IS6 showed significantly lower TMB compared to IS1, IS2, IS3, and IS4, and similar trends were seen with the mutated genes counts as well (Figure 5B). Furthermore, 17 genes including ATM serine/threonine kinase (ATM) were most frequently mutated in each subtype (Figure 5C). These results indicate that the immune subtype can predict TMB and somatic mutation rates in KIRC patients, and that patients with IS1, IS2, IS3, and IS4 may respond positively to mRNA vaccines.

**Figure 5.**
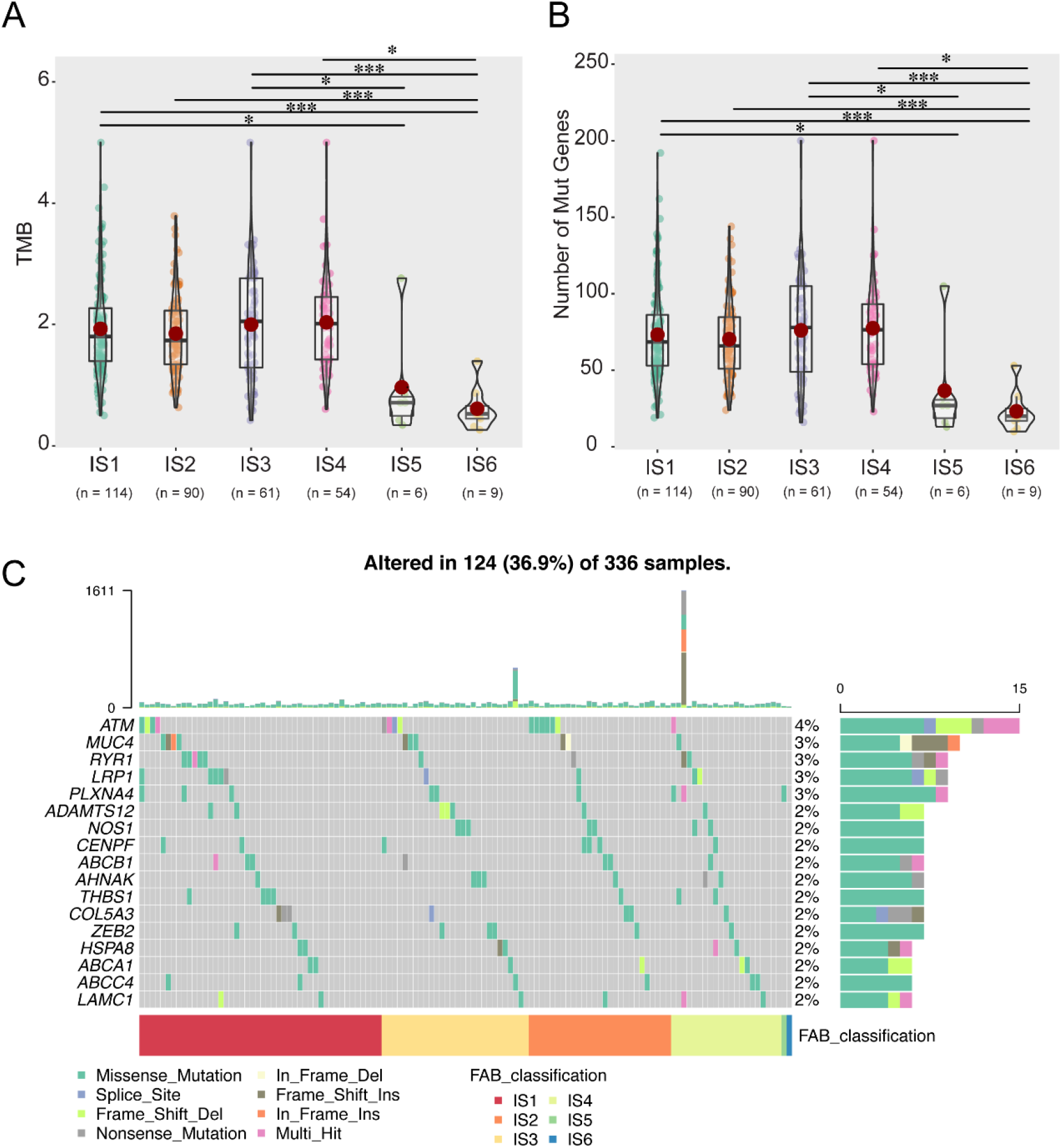
Association between immune subtypes and TMB and mutation. (A) TMB and (B) mutation number in KIRC IS1-IS6. (C) 28 highly mutated genes in KIRC immune subtypes. (\**P* <0.05, ** *P* <0.01, *** *P* < 0.001)

### Association between tumor immune subtypes of KIRC and immune modulators

Current studies have shown that immune checkpoints (ICPs) (e.g., PD-L1, CTLA-4, and TIM-3) and immunogenic cell death (ICD) modulators (e.g., CALR and HMGB1) play important roles in regulating host anti-tumor immunity, thus affecting the efficacy of mRNA vaccines. Therefore, the differential expression of ICPs and ICD modulators in different tumor subtypes of KIRC patients was evaluated. Forty-seven ICPs related genes were detected in TCGA cohort, of which 46 (97.9%) genes except for ICOSLG in TCGA cohort (Figure 6A) were differentially expressed between the immune subtypes. Forty-five ICPs related genes were detected in TCGA cohort, and 44 (97.8%) genes except for TNFSF9 in GEO cohort (Figure 6B) were differentially expressed between the immune subtypes. Such as CD276, CD44, and VTCN1 were significantly upregulated in IS6 tumors in the TCGA cohort, while BTNL2, CD244, CD276, ICOSLG, KIR3DL1, PDCD1, TMIGD2, TNFRSF18, TNFRSF4, TNFRSF8, TNFSF14, TNFSF18, and VTCN1 were overexpressed in the IS6 tumors in GEO cohort.

**Figure 6.**
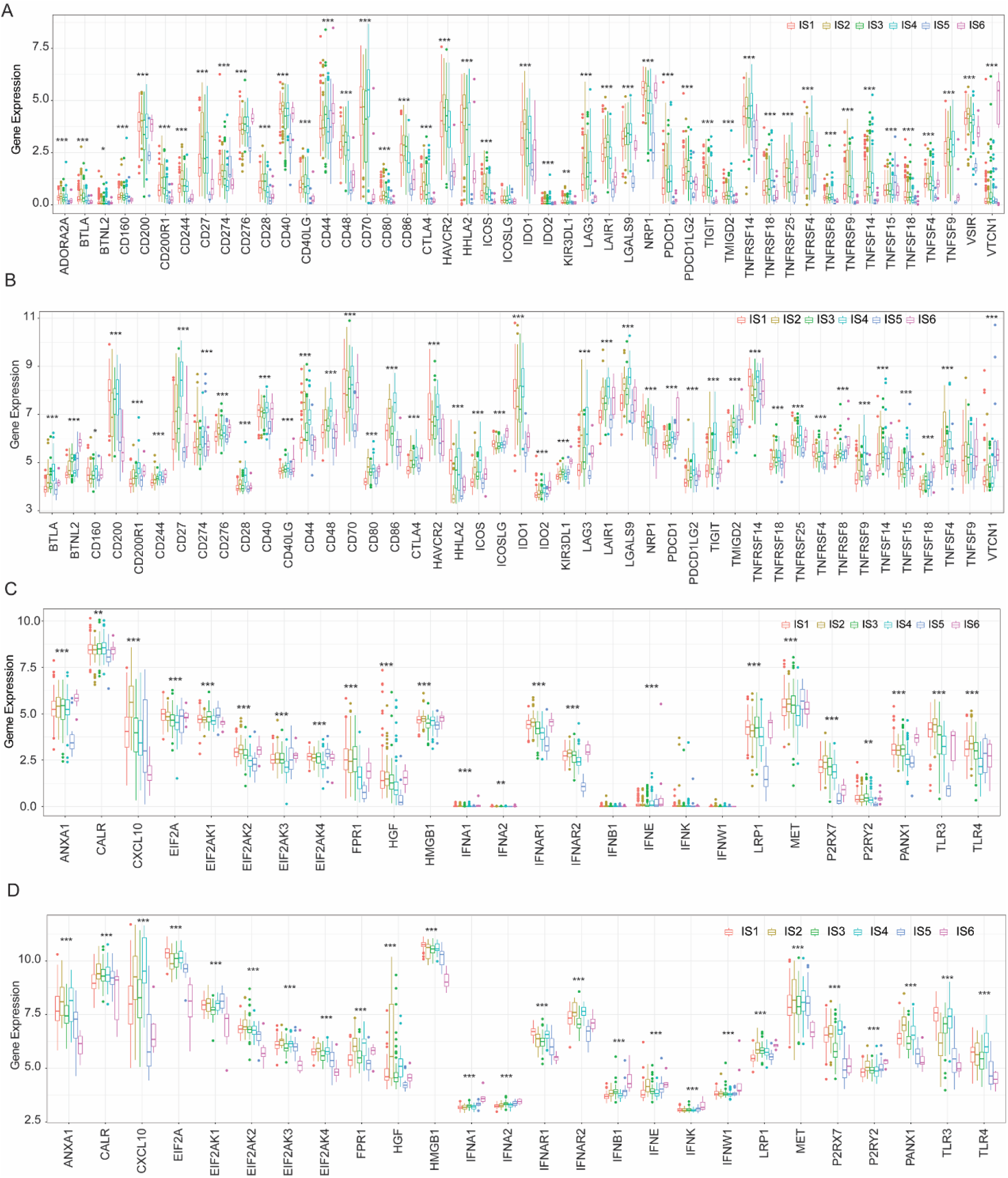
Association between tumor immune subtypes and ICPs and ICD modulators. (A, B) Differential expression of ICP genes among the KIRC immune subtypes in TCGA (A) and GEO (B) cohorts. (C, D) Differential expression of ICD modulator genes among the KIRC immune subtypes in TCGA (C) and GEO (D) cohorts. \**P*<0.05, ** *P* < 0.01, *** *P* < 0.001

Twenty-six ICD genes were detected in TCGA cohort, of which 23 (88.5%) were differentially expressed among the immune subtypes (Figure 6C). Likewise, 26 ICD genes were expressed in GEO cohort, of which 26 (100%) showed significant differences between the subtypes (Figure 6D). For instance, CXCL10, P2RX7, TLR3, and TLR4 were significantly upregulated in IS2 tumors in TCGA cohort, while ANXA1, CXCL10, FPR1, IFNAR2, P2RX7, and TLR4 showed significantly higher expression levels in IS4 tumors in GEO cohort. In conclusion, tumor immunotyping can reflect the expression levels of ICPs and ICD modulators, and may serve as a potential therapeutic biomarker for anti-KIRC mRNA vaccines.

### Cellular and molecular characterization of tumor immune subtypes

The effect of mRNA vaccine on tumor patients depends on the status of tumor immunity. Hence, we further characterized the immune cell components in the 6 immune subtypes by scoring 28 previously reported signature genes in both TCGA and GEO cohorts using ssGSEA. Hence, the immune cell components in 6 immune subtypes were further characterized by ssGSEA with scoring previously reported 28 characteristic genes in TCGA and GEO cohorts. As shown in Figure 7A, the immune cell components were divided into six clusters. IS5 and IS6 showed similar immune cell scores in TCGA cohorts, while similar distribution was observed in IS1-4. More remarkably, the composition of tumor immune cells was consistent across these different subtypes. For instance, the scores of CD56^bright^ natural killer (NK) cells, CD56^dim^ natural killer (NK) cells, eosinophils, immature Dendritic cells, memory B cells, monocytes, neutrophils, plasmacytoid dendritic cells, type 17 T helper cells were significantly higher in IS5 and IS6 compared to IS1-4, (Figure 7B). Thus, IS1-4 are immunological “cold” while IS5 and IS6 are immunological “hot” phenotypes. The results also showed similar trends in the GEO cohort (Figure 7C, D). These results suggest that the immune subtypes reflect the immune status of KIRC and can be used to determine which patients are suitable for mRNA vaccination. The mRNA vaccine produced by these antigens can induce immune infiltration in patients with immunologically “cold” IS1-4 tumors.

**Figure 7.**
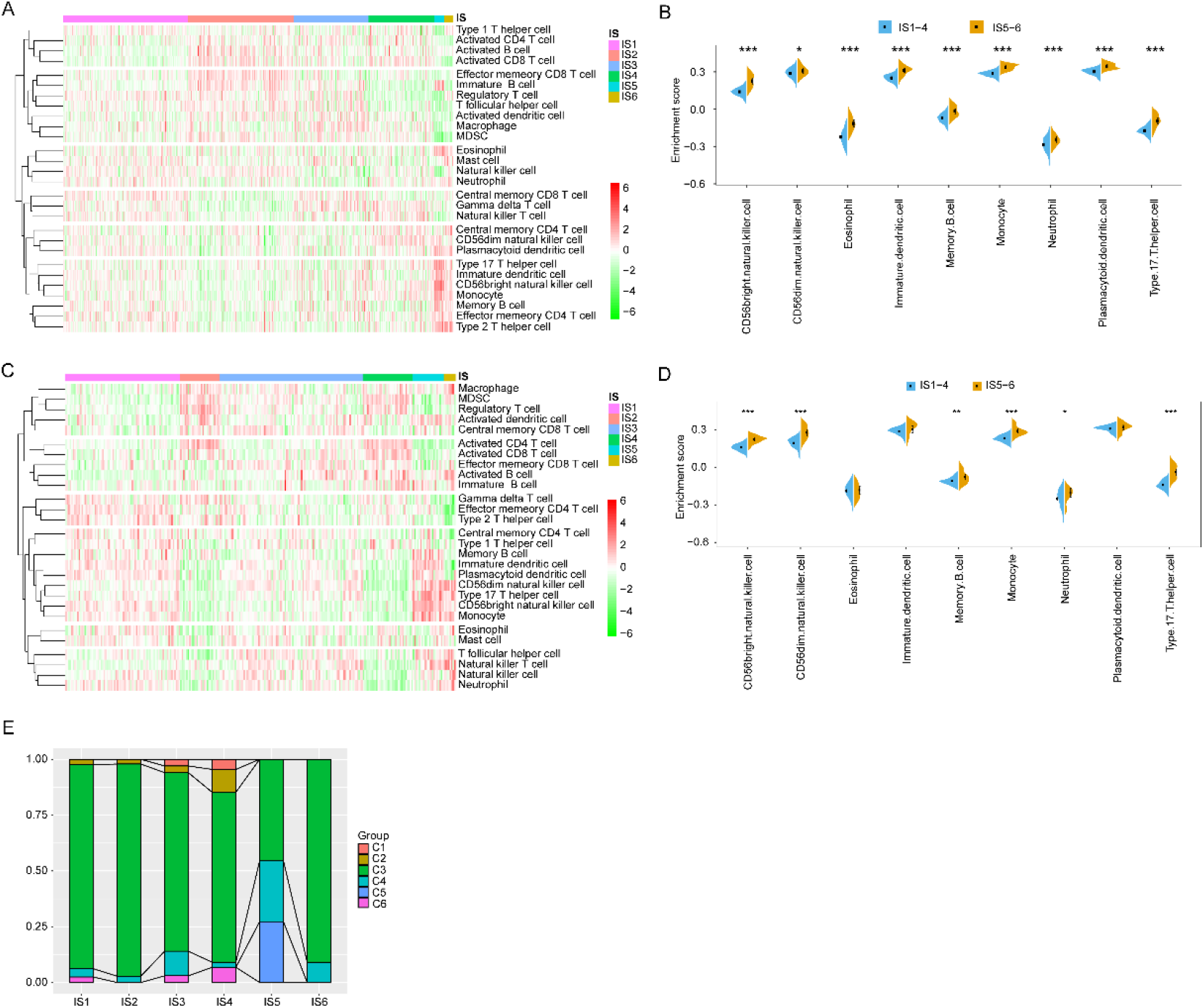
Cellular and molecular characteristics of immune subtypes. (A, C) Differential enrichment scores of 28 immune cell signatures among KIRC immune subtypes in TCGA (A) and GEO (C) cohorts. (B, D) Differential enrichment scores of 9prognostically relevant immune cell signatures in TCGA (B) and GEO (D) cohorts. (E) Overlap of KIRC immune subtypes with 6 pan-cancer immune subtypes.

To demonstrate the reliability of this immunotyping, we next explored the correlation between the 6 immune subtypes and previously reported 6 pan-cancer immune subtypes (C1-C6), of which KIRC was mostly clustered into C3, C4 and C2 [31]. As shown in Figure 7E, IS1, IS2, IS3, IS4, IS6 mainly overlap with C3, IS5 with C3, C4 and C5. C3 and C6 were associated with better and worse prognoses respectively, while C1 and C2 indicated intermediate prognoses. The results showed that patients with IS5 and IS6 tumors had prolonged survival compared with those with IS3 and IS4. Interestingly, the majority of IS5 patients with better prognosis and IS4 patients with the poorest survival overlapped with C3. These findings not only demonstrate the reliability of our immunotyping methods, but also enrich previous classifications. In summary, the immune subtypes reflect the cellular and molecular characteristics of KIRC patients and indicate their immune status. Therefore, immunosubtypes are promising biomarkers for mRNA vaccines, and patients with immunological “cold” tumors of type IS1-4 with or without immunosuppressive microenvironment may be candidates for mRNA vaccines.

### Immune landscape of KIRC

Immune gene expression profiles of individual tumor patients were used to construct the immune landscape of KIRC (Figure 8A). As shown in Figure 8B, the horizonal axis was correlated to various immune cells, of which plasmacytoid dendritic cells, Type 2 T helper cells and memory B cells showed the most positive correlation, while the vertical coordinate was mostly positive associated with effector CD56 bright natural killer cells. The integral distribution of IS2 was opposite to that of IS5 and IS6. In addition, the same subtypes also showed opposite distributions, indicating significant intra-cluster heterogeneity among subtypes, especially IS 1, IS 3, and IS 4. IS2 and IS5 were further divided into two subsets according to their locations of the immune cell population (Figure 8E), and the enrichment scores of immune cells among the multiple subsets were significantly different (Figure 8F). For example, IS2C showed lower counts of activated B cells, activated CD4^+^ T cells, activated CD8^+^ T cells, effector memory CD8^+^ T cells, macrophage, and myeloid-derived suppressor cells (MDSCs), while IS5C scored lower in terms of CD56bright natural killer cell, gamma delta T cell and Type 17 T helper cell. Thus, the mRNA vaccine may be relatively viable in IS5C and more effective in IS2C. In addition, after comparing the prognosis of samples with extreme distributional locations in the immune landscape, the survival probability of patients in group C was the best, which was also consistent with the above results (Figure 8C, D). Taken together, the immune landscape based on immune subtypes can precisely identify immune components of each KIRC patients as well as predict their prognoses, which is favorable for selecting personalized therapeutics for mRNA vaccine. In summary, the immune landscape mapping based on immune subtypes can accurately identify the immune components of each KIRC patient and predict the patient’s prognosis, providing favorable conditions for personalized selection of mRNA vaccines for tumor treatment.

**Figure 8.**
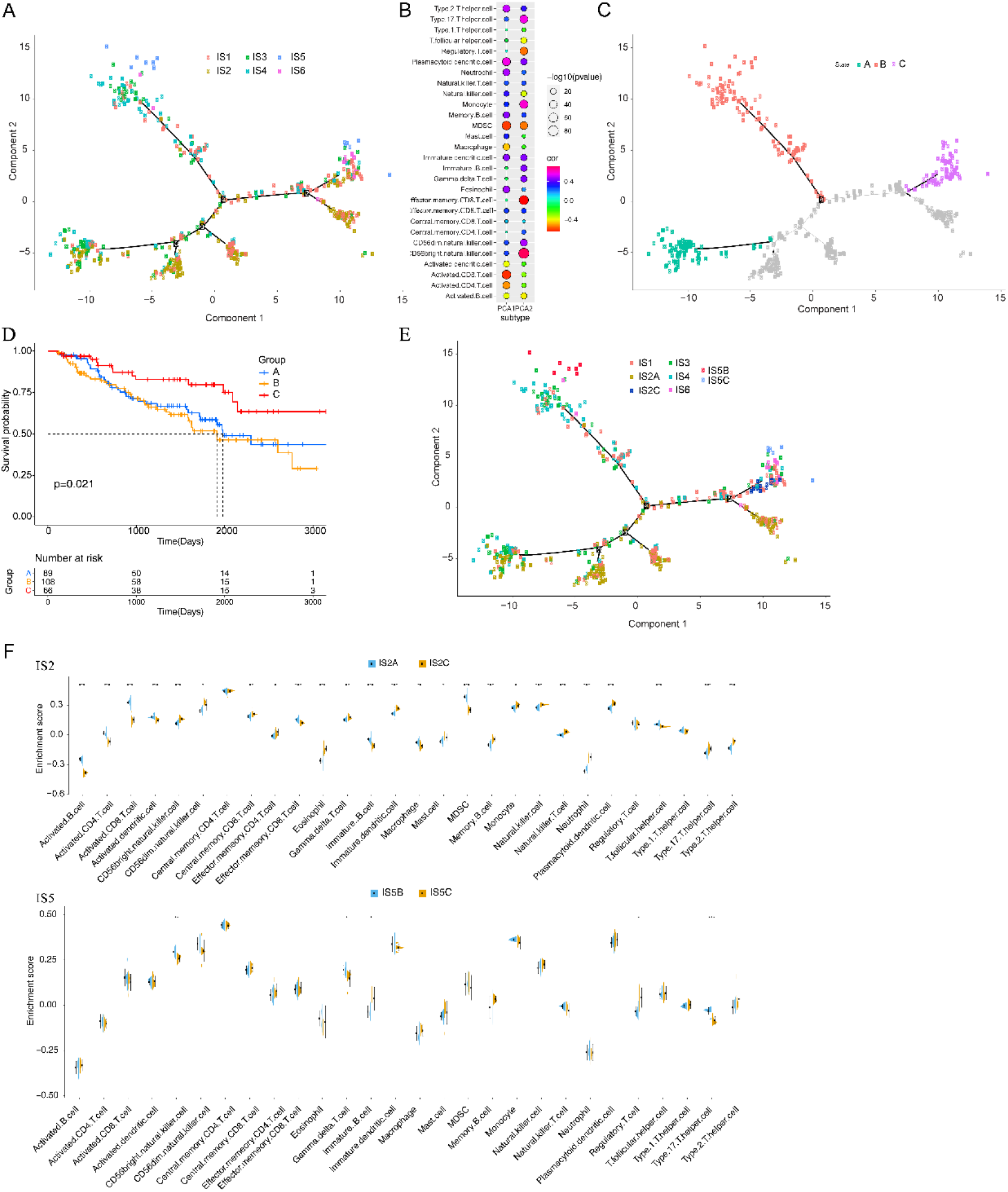
Immune landscape of KIRC. (A) Each point represents a patient, and the immune subtype is color-coded. The horizontal axis represents the first principal component, and the vertical axis represents the second principal component. (B) Heat map of two principal components with 28 immune cell signatures. (C) Patients separated by the immune landscape based on their location. (D) Separated patients were associated with different prognoses. (E) Immune landscape of the subsets of KIRC immune subtypes. (F) Differential enrichment scores of 28 immune cell signatures in the above subsets.

### Identification of immune gene co-expression modules and hub genes of KIRC

Immune gene co-expression modules were identified by clustering the samples by WGCNA (Figure 9A) with a soft threshold of 4 for scale-free network (Figure 9B and C). The representation matrix was then converted to an adjacency and then to a topological matrix. The average linkage hierarchical clustering method was used, with at least 30 genes in each network, in accordance with the standard of hybrid dynamic shear tree. Eigengenes of each module were calculated and the close modules were merged into a new one with height = 0.25, (deep split =2) and min module size = 30. As shown in Figure 9D, 9 co-expression modules with 2108 transcripts were obtained, among which the grey module gene was not clustered with other genes (Figure 9E). After further analyzing the distribution of the 6 immune subtypes in the eigengenes of 8 (except grey) modules, it was found that the distribution of 7 immune subtypes was significantly different (Figure 9F). IS5 showed the lowest eigengenes in red, yellow, blue, and pink modules and IS2 showed the highest eigengenes in the brown, blue and red modules. Further prognostic correlation analysis showed that the brown, yellow, blue, pink, and red modules were significantly associated with the prognosis of KIRC (Figure 10A). Moreover, the brown module was related to T cell receptor signaling pathway, which was positively associated with the component 1 of immune landscape (Figure 10B, C).

**Figure 9.**
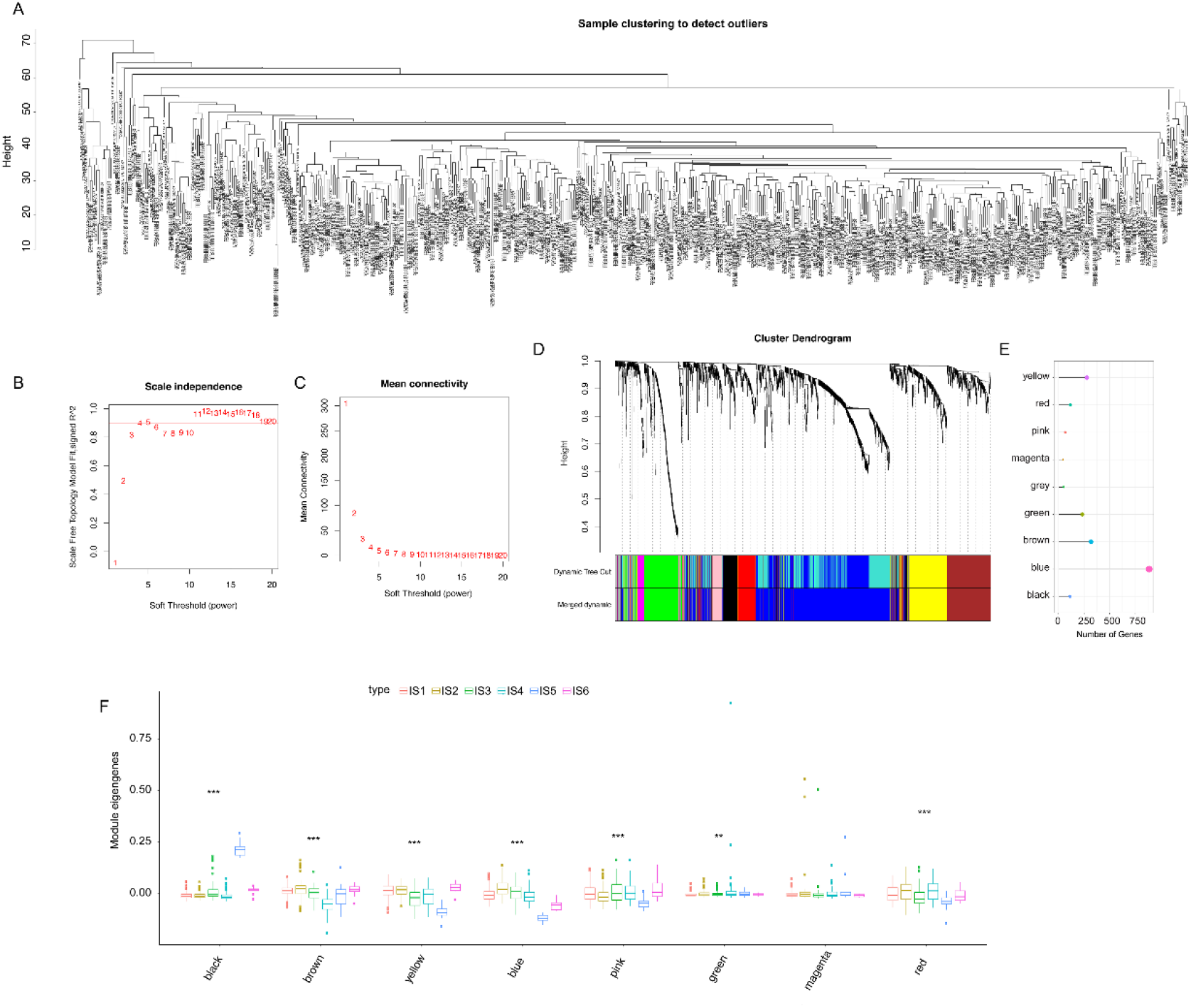
Identification of immune gene co-expression modules of KIRC. (A) Sample clustering. (B) Scale-free fit index for various soft-thresholding powers (β). (C) Mean connectivity for various soft-thresholding powers. (D) Dendrogram of all differentially expressed genes clustered based on a dissimilarity measure (1-TOM). (E) Gene numbers in each module. (F) Differential distribution of feature vectors of each module in KIRC subtypes.

**Figure 10.**
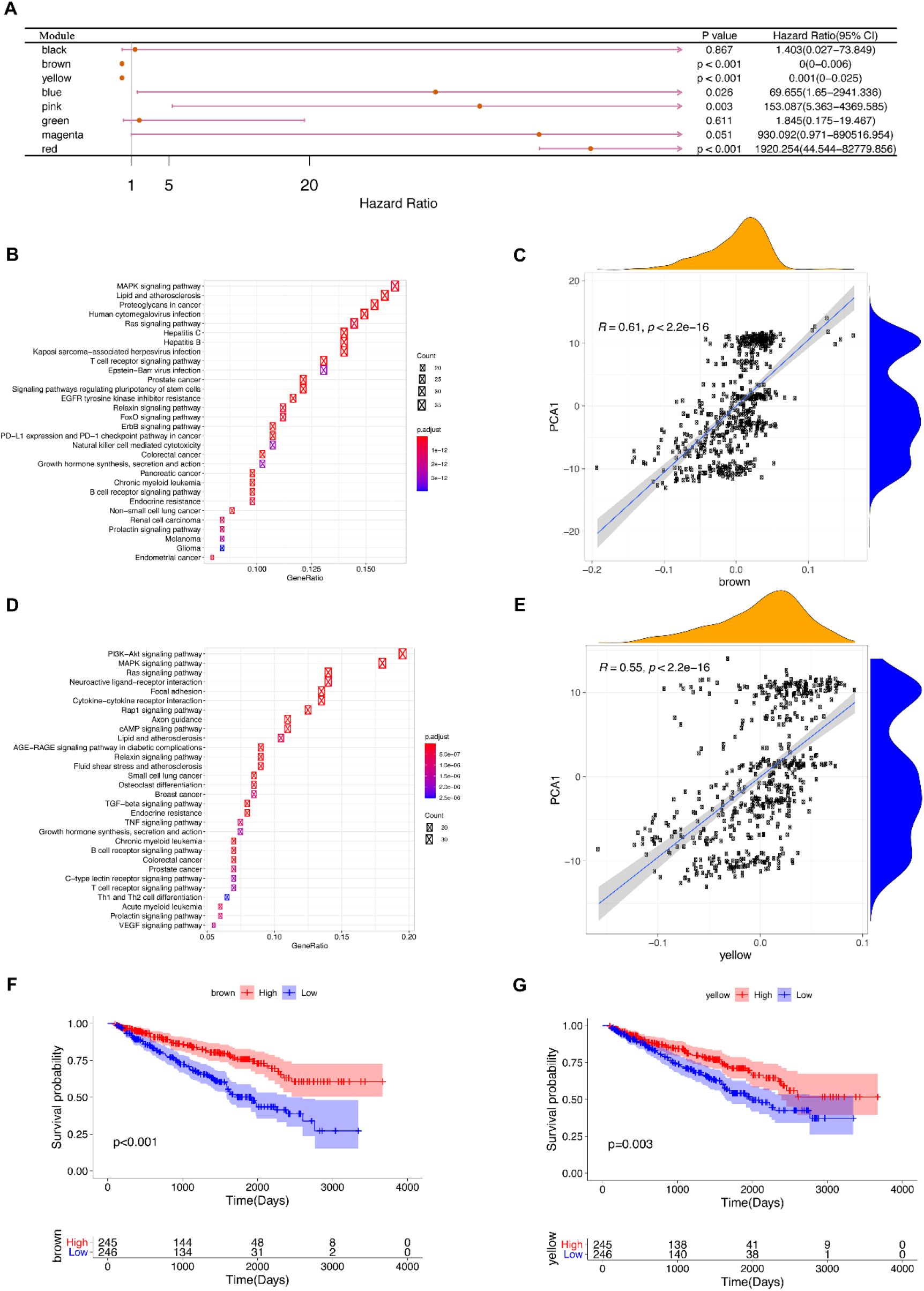
Identification of immune hub genes of KIRC. (A) Forest maps of single factor survival analysis of 8 modules of KIRC. (B) Dot plot showing top 30 KEGG terms in the blue module. The dot size and color intensity represent the gene count and enrichment level respectively. (C) Correlation between blue module feature vector and second principal component in immune landscape. (D) Dot plot showing enriched terms in green module. (E) Correlation between green module feature vector and second principal component in immune landscape. (F) Differential prognosis in brown module with high and low mean. (G) Differential prognosis in yellow module with high and low mean.

Likewise, the yellow module related to T cell receptor signaling pathway showed consistent positive correlation as well (Figure 10D, E). Analysis of prognostically relevant genes of the brown and yellow module showed that higher expression scores correlated with better prognosis in the TCGA cohorts, which is consistent with above mentioned findings (Figure 10F, G). The infiltration and activation of T cells and other immune cells in tumor tissues, as well as inhibition of immune-suppressive cells, largely determine the therapeutic potential of mRNA vaccine in cancer patients with specific immune subtype. Accordingly, mRNA vaccine might be not suitable to patients with high expression of genes clustered into brown and yellow modules. Finally, four hub genes with > 90% relevance in brown and yellow modules were identified, including S1PR1, ETS1, IREB2 and UBR1, which are potential biomarkers for mRNA vaccine.

## Discussion

As far as we are aware, this is the first study to screen KIRC antigens on a large scale based on immune effects for the development of an mRNA vaccine. By constructing the abnormal expression and mutational landscape of KIRC, we identified a series of targeted antigens, including LRP2, and DOCK8, as promising mRNA vaccine candidates. Their upregulation was not only associated with better OS and RFS, but also with higher APC and B cell infiltration. Therefore, these antigens play critical roles in the development and progression of KIRC and can be directly processed and presented to CD8^+^ T cells in the presence of sufficient lymphocyte infiltration to induce an immune attack. Although these candidate genes must be functionally proven to be vaccines, their potential for mRNA vaccine development has been supported by previous reports. For example, Megalin/LRP2 is a receptor that plays an important role in the physiology of multiple organs including kidney, lung, intestine, and gallbladder and in the physiology of the nervous system. LRP2 expression is reduced in fibrosis related diseases, including diabetic nephropathy, liver fibrosis, and cholelithiasis, as well as some breast and prostate cancers [33]. In patients with congenital pulmonary airway malformation, functional destructive mutation of LRP2 were also found to be highly relevant in lung development and cancer [34]. Sustained expression of LRP2 in melanoma cells is critical for cell maintenance, as siRNA-mediated reduction of melanoma cell expression significantly reduces melanoma cell proliferation and survival rates [35]. Statistical analysis also showed that LRP2 was significantly associated with high tumor mutational burden [36]. DOCK8, a gene that encodes a guanine nucleotide exchange factor that is highly expressed in lymphocytes, and mutations in this gene cause an autosomal recessive form of hyperimmunoglobulin E syndrome (AR-HIES), known as DOCK8 deficiency. In humans, DOCK8 deficiency leads to combined immunodeficiency disease (CID), clinically associated with chronic infection of a variety of microbial pathogens, and is prone to malignant tumors [37, 38]. DOCK8 gene and epigenetic inactivation are involved in the development and/or progression of lung cancer and other cancers by interfering with cell migration, morphology, adhesion, and growth [39]. In addition, the high expression of DOCK8 indicates that patients with HPV-positive head and neck squamous cell carcinoma (HNSCC) have a good prognosis and an elevated level of microenvironmental immune infiltration [40].

Given that mRNA vaccine is only beneficial for a fraction of cancer patients, we classified KIRC into six immune subtypes based on immune gene expression profiles for selecting the appropriate population for vaccination. The six immune subtypes exhibited distinct molecular, cellular, and clinical characteristics. Patients with IS5 and IS6 tumors showed better prognosis compared to other subtypes in both GEO and TCGA cohorts, which suggests the immunotyping can be used for predicting the prognoses of KIRC patients. In addition to prognostic prediction, immunotyping is also indicative of the therapeutic response to mRNA vaccine. For instance, patients with IS1 −4 tumors with higher TMB and somatic mutation rates may have greater responsiveness to mRNA vaccine. According to previous immunotyping studies, KIRC was classified into the C1-C6 subtypes. Most patients were clustered into the C3, C4, C2 and C6 subtypes. C3 is associated with superior, C1 and C2 with moderate, and C6 with inferior prognoses [31]. In this study, KIRC was differentiated into IS1-IS6 subtypes. IS1, IS2, IS3, IS4, IS6 mainly overlapped with C3, IS5 with C3, C4 and C5. Studies have shown that C3 and C6 were associated with better and inferior prognoses respectively, while C1 and C2 indicated moderate prognoses. Patients with IS5 and IS6 tumors had prolonged survival compared to those with IS3 and IS4, which is also consistent with previous findings. Interestingly, the majority of IS5 patients with better prognosis and IS4 patients with the lowest survival rate overlapped with C3.

Since the tumor immune status is a determinant of the efficacy of the mRNA vaccine, it is necessary to further characterize the immune cell components of different subtypes [41]. IS5 and IS6 showed significantly elevated scores of CD56bright natural killer cells, CD56dim natural killer cells, eosinophils, immature dendritic cells, memory B cells, monocytes, neutrophils, plasmacytoid dendritic cells and type 17 T helper cells compared to IS1-4. This indicated that IS5 and IS6 are immunological “hot”, and IS1-4 are immunological “cold” phenotypes, which is the lack or paucity of tumor T cell infiltration. Since the presence of immune checkpoints affects the immune response of tumors, ICBs combined with other tumor therapies can induce ICD in tumors, thereby enhancing the anti-tumor effect [42]. The high expressions of ICPs in IS6 tumors in GEO cohort and in TCGA cohort suggest an immunosuppressive tumor microenvironment, which may inhibit the mRNA vaccine from eliciting an effective immune response. In contrast, the elevated expression of ICD modulators in IS2 tumors in TCGA cohort, and in IS4 tumors in GEO cohort are immunological “cold”, which are suggestive of greater potential of mRNA vaccine in these immune subtypes. Interestingly, CXCL10 is also an ICD modulator, and may therefore be a stronger candidate for mRNA vaccine compared to the other selected antigens. In addition, the complex immune profile of KIRC indicates considerable heterogeneity between individual patients and within the same immune subtype, narrowing the immune components available for the development of personalized mRNA vaccine therapies.

The molecular signatures of these tumors were consistent with the immune signatures, indicating that patients with different immune subtypes respond distinctly to mRNA vaccine. For example, IS2C showed lower counts of activated B cells, activated CD4^+^ T cells, activated CD8^+^ T cells, effector memory CD8^+^ T cells, macrophage, and myeloid-derived suppressor cells (MDSCs), indicating an immunologically cold phenotype. Thus, the mRNA vaccine may be relatively viable in IS5C and more effective in IS2C. In order to avoid the low immunogenicity of IS1-4 tumors, the use of an mRNA vaccine to stimulate the immune system by triggering immune cell infiltration may be an appropriate option. For the immune cold and immunosuppressive IS2C tumors exhibiting low CD8^+^ T cells the least lymphocyte, combining the vaccine with immune checkpoint blockage (ICB) or ICD modulators may reinvigorate the immune system and increase immune cell infiltration. S1PR1, ETS1, IREB2, and UBR1 were identified as hub genes in the brown and yellow modules, and their upregulation was positively associated with the component 1 of immune landscape, suggesting that patients expressing high levels of these genes may not respond to the mRNA vaccine. Therefore, the immunotyping method presented in this paper is reliable and a good supplement to the previous classification methods. Nevertheless, the vaccine antigens and other prognostic markers identified in this study need to be further validated in future studies and in preclinical studies.

## Conclusions

Based on a comprehensive analysis of the immune characteristics and prognosis of tumor cells, we suggest that LRP2, and DOCK8 are potential KIRC antigens for mRNA vaccine development, and patients with immune subtypes IS1-4 are suitable for vaccination. These results provide a theoretical basis for the development of anti-KIRC mRNA vaccine, prediction of patient prognosis and selection of appropriate patients for vaccination.

## Abbreviations

APC: Antigen presenting cells
DCs: Dendritic cells
DOCK8: dedicator of cytokinesis 8
ICB: Immune checkpoint blockage
ICD: Immunogenic cell death
ICPs: Immune checkpoints
IVT: in vitro transcribed
KIRC: Kidney renal clear cell carcinoma
LRP2: LDL receptor related protein 2
MET: MET proto-oncogene receptor tyrosine kinase
MDSCs: Myeloid-derived suppressor cells
OS: Overall survival
RCC: Renal cell carcinoma
RFS: Relapse-free survival
ssGSEA: Single-sample GSEA
TCGA: The Cancer Genome Atlas
TIICs: Tumor infiltrating immune cells
TIME: Tumor immune microenvironment
TMB: Tumor mutation burden

## Acknowledgements

The authors would like to thank GEO (https://www.ncbi.nlm.nih.gov/geo) and TCGA (https://www.cancer.gov/tcga) for data collection, as well as GEPIA2 (http://gepia2.cancer-pku.cn), cBioPortal (http://www.cbioportal.org, version v3.2.11) and TIMER (https://cistrome.shinyapps.io/timer) for the provision of data processing and customizable functions.

## Authors’ contributions

S.C.Z., Y.O.Y. and J.P. designed the study, S.C.Z. and Y.X. performed the bioinformatics analysis and prepared the figures; S.K., C.M., and Y.W. participated in data collection and analysis. S.C.Z., Z.Z., J.P., and Y.O.Y. interpreted the results, wrote, and revised the manuscript. All authors read and approved the final manuscript.

## Availability of data and materials

All data generated and described in this article is available from the appropriate Web server and is freely available to any scientist who wishes to use it for non-commercial purposes, without violating the confidentiality of the participants. Further information is available at the reasonable request of the corresponding author.

## Ethics approval and consent to participate

Not applicable.

## Consent for publication

Not applicable.

